# Does researcher activity affect *Dromiciops gliroides* capture rates? A test in two contrasting forest habitats

**DOI:** 10.1101/801332

**Authors:** Daniela A. Salazar, Francisco E. Fontúrbel

## Abstract

Aiming to understand the low *D. gliroides* capture rates at the Valdivian Coastal Reserve, we disposed camera-traps at two contrasting forest habitats: a native forest habitat and a transformed habitat composed by a *Eucalyptus globulus* plantation with native understory vegetation. Camera-trap survey was conducted before and during live-trap operation. We found a large number of photographic records at the pre-bait period (41 photos at the native habitat and 22 at the transformed habitat). Then, when we conducted the live-trapping survey, photographic records decrease to 7 at the native habitat and 5 at the transformed habitat. Compared to similar locations in southern Chile, our study site shows low sampling effectiveness and capture success values, which could be explained by the disturbance generated by the researcher by checking the trap grid in a daily basis in a remote place where human presence is sporadic.

## Introduction

The *monito del monte* (*Dromiciops gliroides* Thomas 1894) is an arboreal marsupial endemic to the temperate rainforests of South America (36 – 43 °S). This species has a keystone ecological role as seed disperser at these forests (Amico and Aizen, 2000; Amico et al., 2009), and its conservation is a priority as it is the only extant species of the Order Microbiotheria. This species was considered to be rare because of the low abundances reported when small mammals are assessed with traditional methods (Kelt, 2000; Meserve et al., 1988; Patterson et al., 1989) but with the new sampling method (Fontúrbel and Jiménez, 2009) yielded capture rates of up to 11 %. Consequently, the previously reported low abundances may result from a sampling artifact.

The current knowledge on the ecological aspects of *D. gliroides* was synthesized by Fontúrbel et al. (2012), comparing the available information from Argentina and Chile, finding no significant differences in capture rates between both countries. However, in our study site, the Valdivian Coastal Reserve, the capture rates were much lower than those previously reported (0.38 % during the 2011-2012 austral summer; unpublished data). Given this, we installed camera-traps in front of the live traps at two contrasting habitats, aiming to assess whether the marsupial was in fact less abundant compared to other locations, or the differences were due to problems in the sampling method. The large number of photographic records obtained at both habitats in a previous study (Fontúrbel et al., 2014), suggests that this species is more abundant than we initially thought, but for an unknown reason it is not being captured.

A topic rarely addressed in field ecology studies is the potential effects of human presence on capture rates. Studies conducted on fear ecology are mainly focused on large mammals and are conducted in controlled conditions (Clinchy et al., 2013; Ripple and Beschta, 2004). Nevertheless, little is known about the potential responses of wild small mammals to human presence and sporadic habitat disturbance.

We hypothesized that the researcher activity would affect *D. gliroides* capture rates at the Valdivian Coastal Reserve because this is a little disturbed and remote area, with very low daily human impact, compared to other known populations of the monito del monte. Consequently, the daily disturbance to which grids are submitted when checked, it would explain at least in part the decrease in capture rates. We evaluated this in two contrasting habitats: an old-growth native forest stand, and a non-managed *Eucalyptus globulus* Labill. plantation with native understory vegetation.

## Methods

The study was conducted at the Valdivian Coastal Reserve (39°57’S, 73°34’W), a 50,530-ha private protected area located in southern Chile, and managed by The Nature Conservancy (Delgado, 2010). This Reserve represents a large continuum with a complex habitat mosaic comprising old- and second-growth native stands (regenerated after clear-cutting), and exotic *Eucalyptus globulus* plantations (12-20 years old, currently unmanaged) with abundant native understory vegetation.

Aiming to capture *Dromiciops gliroides*, we established live-trapping grids in the two contrasting forest habitats during the austral summer of 2014: one grid was set at an old-growth native stand, and the other one at the *Eucalyptus globulus* plantation with native understory. We set 48 Tomahawk-like traps (13×13×26 cm, custom made), disposed in a 6×8 trap lattice, in each trapping grid. Traps were separated 10 m from each other and baited with fresh banana slices (Fontúrbel and Jiménez, 2009; Fontúrbel et al., 2010). Live trapping met the guidelines of the American Society of Mammalogists (Sikes et al., 2011), and was authorized by the Chilean Agriculture and Livestock Bureau (SAG; resolution 8291/2013).

At each trapping grid, we randomly selected ten trapping locations to install camera-traps (Bushnell Trophy Cam 2011) in front of the traps. Camera-trap monitoring was divided in two seven-day periods: (1) a pre-bait period from January 21 to January 28, in which traps were closed and bait was placed above the trap to attract *D. gliroides*; and (2) an operation period from February 10 to February17, in which traps were baited and checked daily early in the morning (bait was renewed every 48 hours). Weather and resource availability conditions between January and February were similar (Fontúrbel, 2013), and the activity patterns of *D. gliroides* were also similar in the two previous summer seasons (Fontúrbel et al., 2014).

The effective sampling area of a grid (camera-trap based or live-trap based) is usually estimated by the grid area plus a buffer of half the largest recapture distance (following Parmenter et al., 2003). However, as we did not recapture any individual, we used a more conservative approach using half of the distance between traps (*i.e.*, 5 m) as buffer distance, which gave us an area of 0.48 ha for each grid.

Camera traps were set in photographic mode with an inactivity interval of 60 s to avoid repetitive shots of the same individual. As we were unable to distinguish individuals from each other (*i.e.*, records may not be independent from each other), we considered our data as relative activity instead of abundance. Aiming to compare the sampling effectiveness between different sites where *D. gliroides* were captured, we estimated the sampling effectiveness, defined as the percentage of individuals actually captured related to those that could be captured. Such expected number was estimated considering the average density and the extent of the grid. For an average density of 23 individual × ha^-1^ and a home range overlap of 50 % (2012; Fontúrbel et al., 2010), the number of individual that are likely to be found in a 0.48 ha grid is: Individuals = 23 individuals/ha × 0.48 ha × (1/0.5) = 22.08 individuals (rounded to 22). Capture succes was estimated as the number of captures obtained, divided by the number of trap-nights disposed and the operation in each grid.

## Results

At both habitats, the largest photographic record was obtained for the pre-bait period, with a total of 63 photos of *D. gliroides*, while for the operation period we obtained only 12, obtaining a large number of records in *Eucalyptus* plantation habitat than in the native forest (46 and 29, respectively including pre-bait and operation periods; figure 1). Examining those figures by day, in both habitats most records were obtained at the second and third day after the beginning of both pre-bait and operation periods (figure 2). In the native habitat, the total number of captures was seven. Considering that they should have been about 22 individuals, the sampling effectiveness was 31.8 %, while the capture success (with seven trap-nights) was 2.1 %. For transformed habitat we obtained five captures, resulting in a sampling effectiveness of 22.7 %, and a capture succes (with 5 trap-nigths) of 2.1 %. For both habitat, we found no correlation between the number of live captures and the number of photographic records during the operation period (Spearman rank correlation, r_s_ = 0.21, *P* = 0.65 for the native habitat; r_s_ = 0.08, *P* = 0.87 for the transformed habitat).

**Figure 1.**
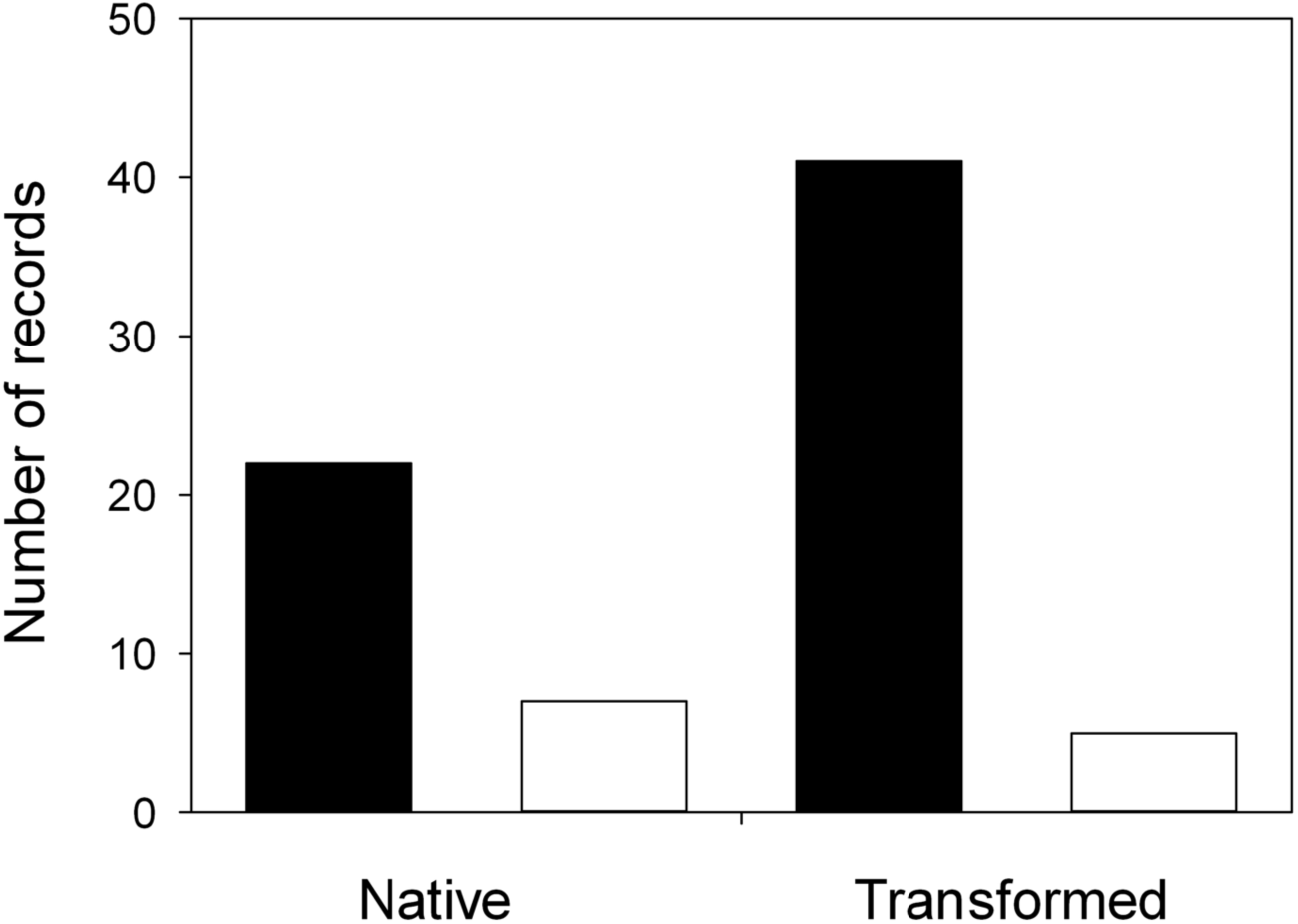
Number of photographic records obtained at each habitat, black bars represent the native forest habitat and white bars represent the *Eucalyptus* plantation habitat.

**Figure 2.**
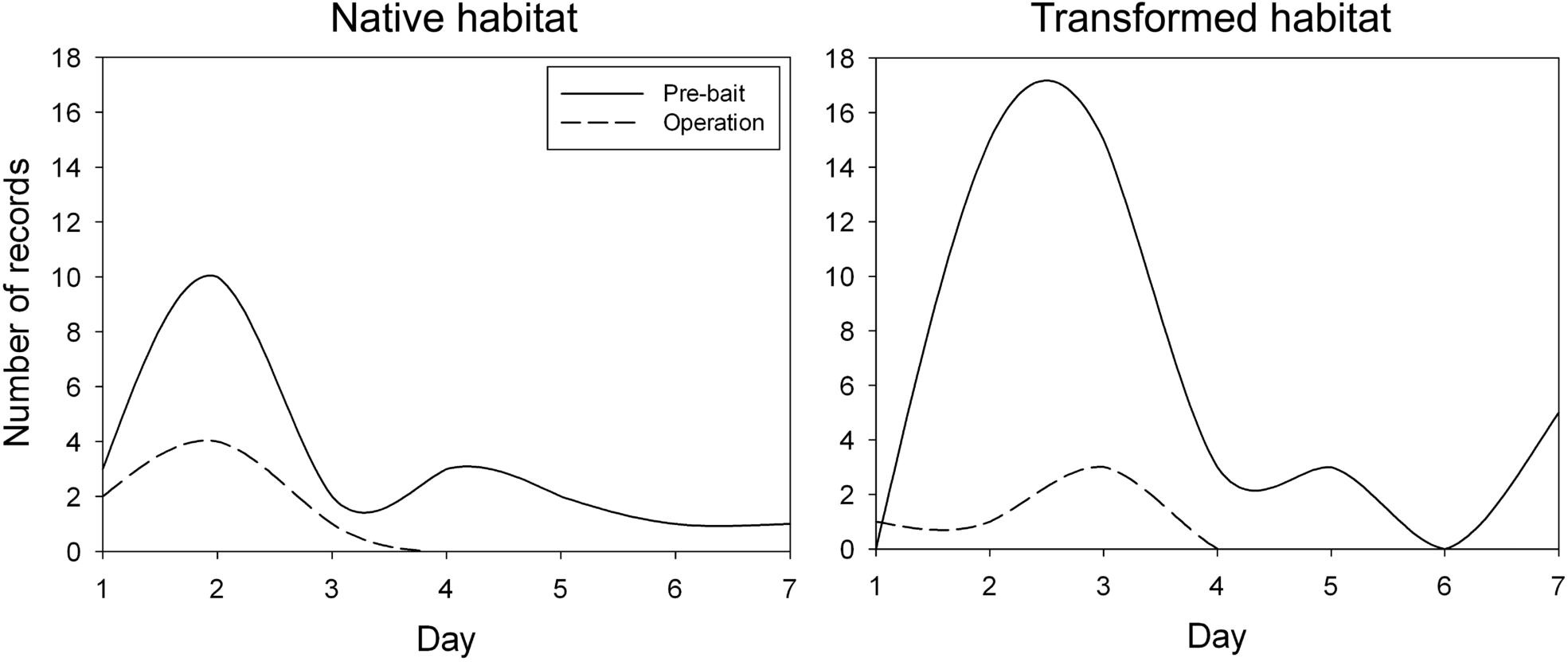
Daily photographic records taken during pre-bait and operation period at native and transformed habitats.

## Discussion

Considering that our study site is a private protected area, it has a much less disturbance and human activity than other locations where *D. gliroides* has been previously captured (Fontúrbel et al., 2012). Consequently, in this site human presence effects are expected to be much higher than in other sites with constant human visitation such as Cascadas and San Martín in Chile, and Llao-Llao in Argentina, which receive many tourists and researchers in a regular basis during the whole summer season, and hence *D. gliroides* individuals may be habituated to human presence. In our study site, however, sporadic disturbance could be reflected in low capture rates, being this effect stronger at the transformed habitat, which is more exposed to predators such as feral dogs and the Rufous-legged owl (*Strix rufipes*; unpublished data).

Comparing the capture success and the sampling effectiveness from Cascadas (comparable in terms of structure, species composition, and resource offer to our native habitat site) and our study site, fewer individuals were captured in our study site than in Cascadas (capture success, Cascadas: 17 %, our site: 2.1 %; sampling effectiveness, Cascadas 50 %, our site: 31.8 %). The use of camera traps in our study arises the possibility that this low capture rates should not be necessarily related to the absence or low abundances of *D. gliroides*, as we initially supposed. This idea is supported by our results, because we obtained 63 camera trap records during the pre-bait period. This shows that despite the marsupial is present in the area, captures are lower during the operation period, this could be explained by the disturbance generated by the researcher visiting the sites to check the traps and replace the bait on a daily basis. Live capturing might be influencing the decrease in the number of photographic records as captured individuals cannot the further photographed, however we found no correlation between the number of captures and photographic records at both habitats. Therefore, we propose that it is necessary to use complementary methods such as camera-traps and pre-baiting, in order to determine whether the absence of *D. gliroides* captures results from a real absence or from a stressful situation imposed by the researcher. If the second case is true, we recommend minimizing the disturbance and reduce the work time in the forest as much as possible. If further data is needed, we recommend gathering it in a different time than the live trapping.

## Acknowledgements

The Rufford Small Grants Foundation (grant 14669-2) and the FONDECYT Project 3140528 (to FEF) funded this study. We are grateful to The Nature Conservancy and the Valdivian Coastal Reserve for granting access permissions and providing field facilities. Comments of C. González-Browne and C. Botto-Mahan improved an earlier version of this manuscript.

## References

Amico, G.C., Aizen, M.A., 2000. Mistletoe seed dispersal by a marsupial. Nature 408: 729–730.

Amico, G.C., Rodríguez-Cabal, M.A., Aizen, M.A., 2009. The potential key seed-dispersing role of the arboreal marsupial *Dromiciops gliroides*. Acta Oecologica 35: 8–13.

Clinchy, M., Sheriff, M.J., Zanette, L.Y., 2013. Predator induced stress and the ecology of fear. Functional Ecology 27: 56–65.

Delgado, C., 2010. Plan de Manejo de la Reserva Costera Valdiviana The Nature Conservancy, Arlington VA.

Fontúrbel, F.E., 2013. Effects of habitat transformation on the ecoevolutionary dynamics of plant-animal mutualisms on a hemiparasitic mistletoe, Facultad de Ciencias. Universidad de Chile, Santiago de Chile, p. 144.

Fontúrbel, F.E., Candia, A.B., Botto-Mahan, C., 2014. Nocturnal activity patterns of the monito del monte (*Dromiciops gliroides*) in native and exotic habitats. Journal of Mammalogy 95: 1199–1206.

Fontúrbel, F.E., Franco, M., Rodríguez-Cabal, M.A., Rivarola, M.D., Amico, G.C., 2012. Ecological consistency across space: a synthesis of the ecological aspects of *Dromiciops gliroides* in Argentina and Chile. Naturwissenschaften 99: 873–881.

Fontúrbel, F.E., Jiménez, J.E., 2009. Underestimation of abundances of the monito del monte (Dromiciops gliroides) due to a sampling artifact Journal of Mammalogy 90: 1357–1362.

Fontúrbel, F.E., Silva-Rodríguez, E.A., Cárdenas, N.H., Jiménez, J.E., 2010. Spatial ecology of monito del monte (*Dromiciops gliroides*) in a fragmented landscape of southern Chile. Mammalian Biology 75: 1–9.

Kelt, D.A., 2000. Small mammal communities in rainforest fragments in central southern Chile. Biological Conservation 92: 345–358.

Meserve, P.L., Lang, B.K., Patterson, B.D., 1988. Trophic relationships of small mammals in a Chilean temperate rainforest. Journal of Mammalogy 69: 721–730.

Parmenter, R.R., Yates, T.L., Anderson, D.R., Burnham, K.P., Dunnum, J.L., Franklin, A.B., Friggens, M.T., Lubow, B.C., Miller, M., Olson, G.S., 2003. Small-mammal density estimation: a field comparison of grid-based vs. web-based density estimators. Ecological monographs 73: 1–26.

Patterson, B.D., Meserve, P.L., Lang, B.K., 1989. Distribution and abundance of small mammals along an elevational transect in temperate rainforests of Chile. Journal of Mammalogy 70: 67–78.

Ripple, W.J., Beschta, R.L., 2004. Wolves and the ecology of fear: can predation risk structure ecosystems? BioScience 54: 755–766.

Sikes, R.S., Gannon, W.L., Care and Use Committee, 2011. Guidelines of the American Society of Mammalogists for the use of wild mammals in research. Journal of Mammalogy 92: 235–253.

